# SNT: A Unifying Toolbox for Quantification of Neuronal Anatomy

**DOI:** 10.1101/2020.07.13.179325

**Authors:** Cameron Arshadi, Ulrik Günther, Mark Eddison, Kyle I. S. Harrington, Tiago A. Ferreira

## Abstract

Quantification of neuronal morphology is essential for understanding neuronal connectivity and many software tools have been developed for neuronal reconstruction and morphometry. However, such tools remain domain-specific, tethered to specific imaging modalities, and were not designed to accommodate the rich metadata generated by recent whole-brain cellular connectomics. To address these limitations, we created *SNT*: a unifying framework for neuronal morphometry and analysis of single-cell connectomics for the widely used Fiji and ImageJ platforms.

We demonstrate that SNT can be used to tackle important problems in contemporary neuroscience, validate its utility, and illustrate how it establishes an end-to-end platform for tracing, proof-editing, visualization, quantification, and modeling of neuroanatomy.

With an open and scriptable architecture, a large user base, and thorough community-based documentation, SNT is an accessible and scalable resource for the broad neuroscience community that synergizes well with existing software.

---

Quantification of neuronal anatomy is essential for mapping information flow in the brain and classification of cell types in the central nervous system. Although digital reconstruction (“tracing”) of the tree-like structures of neurons —axons and dendrites— remains a laborious task, recent improvements in labeling and imaging techniques allow faster and more efficient reconstructions, with neuroscientists sharing more than 140,000^1^ reconstructed cells across several databases. Powerful toolboxes have been developed for neuronal morphometry (**Sup. Information**). However, such tools can be tethered to specific imaging modalities or remain specialized on specific aspects of neuroanatomy workflows. To address these limitations we established a unifying framework for neuronal morphometry and analysis of single-cell connectomics for the widely used Fiji and ImageJ platforms^2,3^.

We re-invented the popular Simple Neurite Tracer program^4^ to create an open-source, end-to-end solution for semi-automated tracing, visualization, quantitative analyses and modeling of neuronal morphology. All aspects of our software, named *SNT —*Simple Neurite Tracer’s popular moniker— can be controlled from a user-friendly graphical interface or programmatically, using a wide variety of scripting languages (**Fig. 1a**).

**Figure 1.**
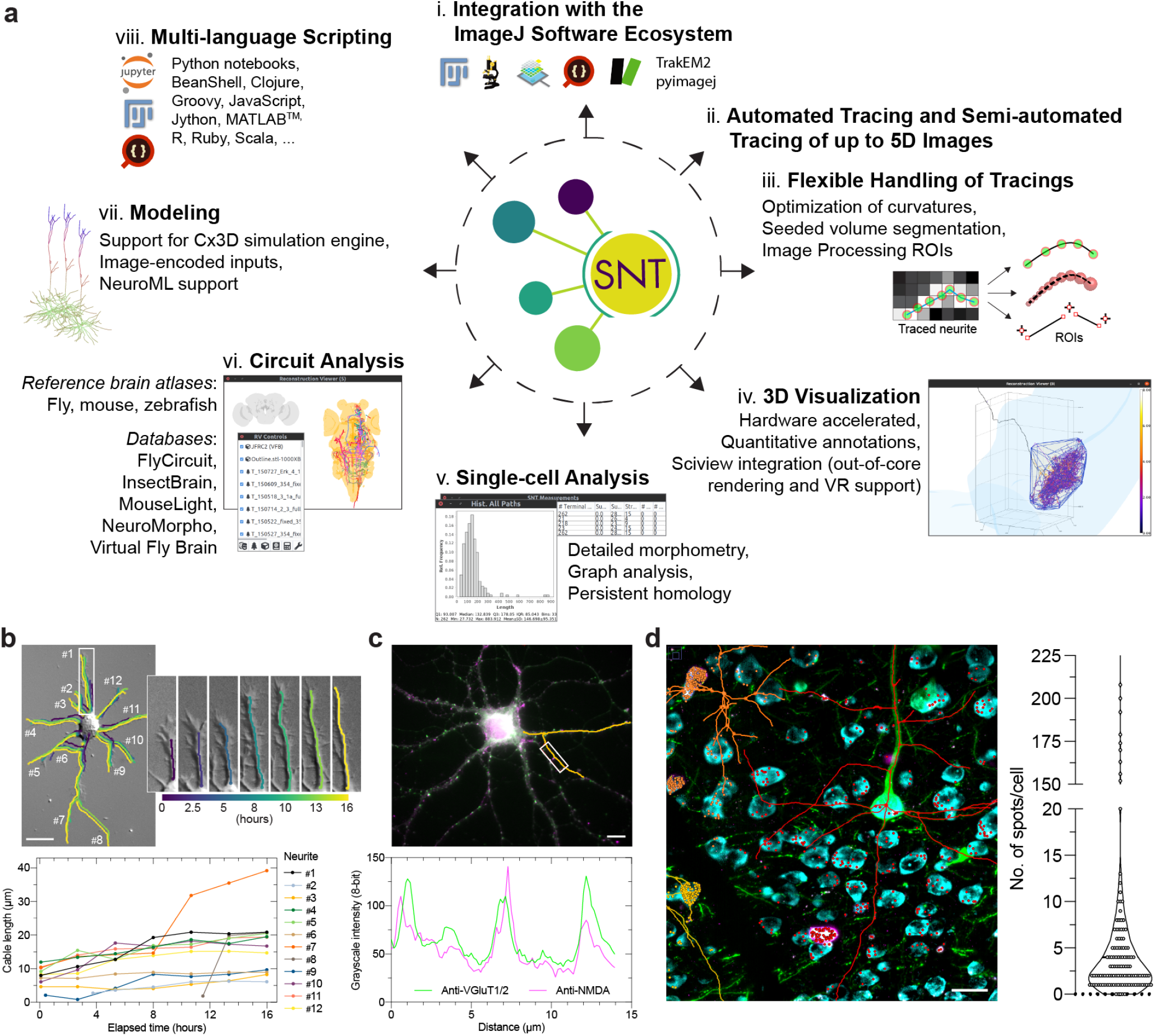
SNT as an end-to-end platform for data retrieval, visualization, quantification, and modeling of neuroanatomical data. (**a**) **Schematic diagram of the software.** (clockwise): i) SNT is powered by the stack of ImageJ-based software, including: Fiji, ImageJ2, sciview, SciJava, ImgLib2, TrakEM2 and pyimagej. ii) Reconstructions can be obtained directly from thresholded images or using semi-automated procedures that support time-lapse and multi-channel light-microscopy imagery. iii) Once center-line reconstructions (“tracings”) are obtained, they can be conveniently processed in subsequent image processing routines. iv) Dedicated neuroanatomy viewers allow for effective quantitative visualization of complex data. v) In addition to single-cell morphometry, vi) circuit analyses are facilitated through support of several online databases and reference brain atlases (Drosophila, mouse and zebrafish). vii) Biophysical modeling of neuronal growth is achieved through Cx3D integration (Sup. Information). viii) Users may use SNT has a standalone interactive program or as a multi-language scripting library. (**b—d) ImageJ interoperability allows for complex data retrieval.** (**b**) Static frame from a non-fluorescent time-lapse video monitoring the development of neuron polarity in a hippocampal neuron growing *in vitro*^19^. The 12 highlighted neurites were traced throughout the video sequence and color coded across time as per color ramp. Insert details growth of neurite #1 at selected time-points. Plot depicts growth dynamics of individual neurites across time. (**c**) Multichannel image of a hippocampal neuron stained *in vitro* for the presynatic markers VGluT1-2 (green) and the postsynaptic NMDA receptor (magenta)^18^. Dendrites were traced (orange) and intensity profiles obtained directly from the tracings. Profiled maxima from the marked region depict synaptic locations. (**d**) Maximum intensity projection of a three-channel 3D image of a gelled-brain section processed for expansion fluorescence in situ hybridization (ExFISH). Dendrites of GFP-labeled neurons (green) were traced in SNT (center-lines for three cells displayed in orange, red, and yellow). Foci reporting on Somatostatin (SST) mRNA (magenta) were detected on neighboring somata, segmented from a counterstain for total RNA (cyan). Point ROIs reporting foci (circles) were labeled with the same hue of the closest traced cell. All procedures performed within ImageJ. Right: Violin plot of SST expression for segmented cells in the sub-volume (N=147). Scale bars: (b,c): 10μm; (d): 20μm.

For semi-automated tracing we implemented a host of new features (described in Sup. Information), including support for multi-channel, and time-lapse images, optimized search algorithms and image processing routines that better detect neuronal processes, and made possible to reconstruct simple morphologies directly from thresholded images. With timelapse sequences, traced paths can be automatically matched across frames so that growth dynamics of individual neurites can be monitored across time. (**Fig. 1b**). To expedite the proof-editing of traced structures, SNT allows users to edit, tag, sort, filter, and rank traced segments, using either ad-hoc labels or morphometric traits. Altogether, these features improve reconstruction accuracy and tracing efficiency.

For visualization, SNT features an interactive 3D viewer dedicated to neuron morphology —*Reconstruction Viewer—* that is hardware accelerated, supports rendering of meshes and detailed annotation of morphometry data. In addition, SNT also integrates with sciview^5^, a visualization tool for mesh-based data and arbitrarily large image volumes, supporting virtual, and augmented reality (**Sup. Information**).

For data retrieval, SNT provides seamless integration with the ImageJ platform, and thus tracing and reconstruction analyses can be intermingled with image processing workflows. To exemplify this, we used SNT to quantify challenging datasets: expression of synaptic markers along dendrites (**Fig. 1c**), and fluorescent in situ hybridization (FISH) imaging of mRNA in the same volume in which dendrites of pyramidal cells were reconstructed (**Fig. 1d**).

Another strength of SNT is that it can connect directly to all the major neuroanatomy databases, including FlyCircuit^41^, InsectBrain^6^, MouseLight^7^, NeuroMorpho^1^, and Virtual Fly Brain^8^, supporting several multi-species brain atlases (Drosophila, mouse, zebrafish, **Fig. 1a**, **Sup. Information**). While such online databases are highly queryable, they remain constrained by website design limitations. Scripting frameworks that can programmatically parse their data bypass those restrictions, facilitate data sharing, scientific reproducibility, bridge isolated data repositories, and promote the development of new tools and features. Importantly, since SNT adopts the SciJava framework^9,10^, it can be scripted using popular computer languages such as Python (and Jupyter notebooks) through pyimagej, Clojure, Groovy, JavaScript, Jython, MATLAB™, R, Ruby, or Scala.

To demonstrate the analytical power of SNT, we parsed the MouseLight database, currently containing the most complex reconstructions described in the literature^7^. In particular, we focused on quantifying the repertoire of strategies adopted by individual cells to broadcast information across the brain. Since no synaptic strengths are currently known for MouseLight neurons, projection strength to target areas must be inferred from morphometric surrogates. In a programmatic, unbiased approach, we used two morphological criteria (normalized cable length and number of axonal endings) to retrieve the number of anatomical brain areas innervated by individual axons (**Fig. 2a**). In doing so, we identified two extremes of connectivity: cells that connect exclusively to a single projection brain area, and cells that project broadly over a multitude of brain areas (**Fig. 2b**). A key feature of SNT is the ability to generate streamlined connectivity diagrams, holding quantitative information determined from the intersection or union of multiple morphometric criteria that can be customized using SNT’s interactive tool *Graph Viewer.* These type of diagrams can be generated at the single-cell level (**Fig. 2c**), or from cell populations (**Fig. S7**), and are a valuable visualization tool for connectomics^11^.

**Figure 2.**
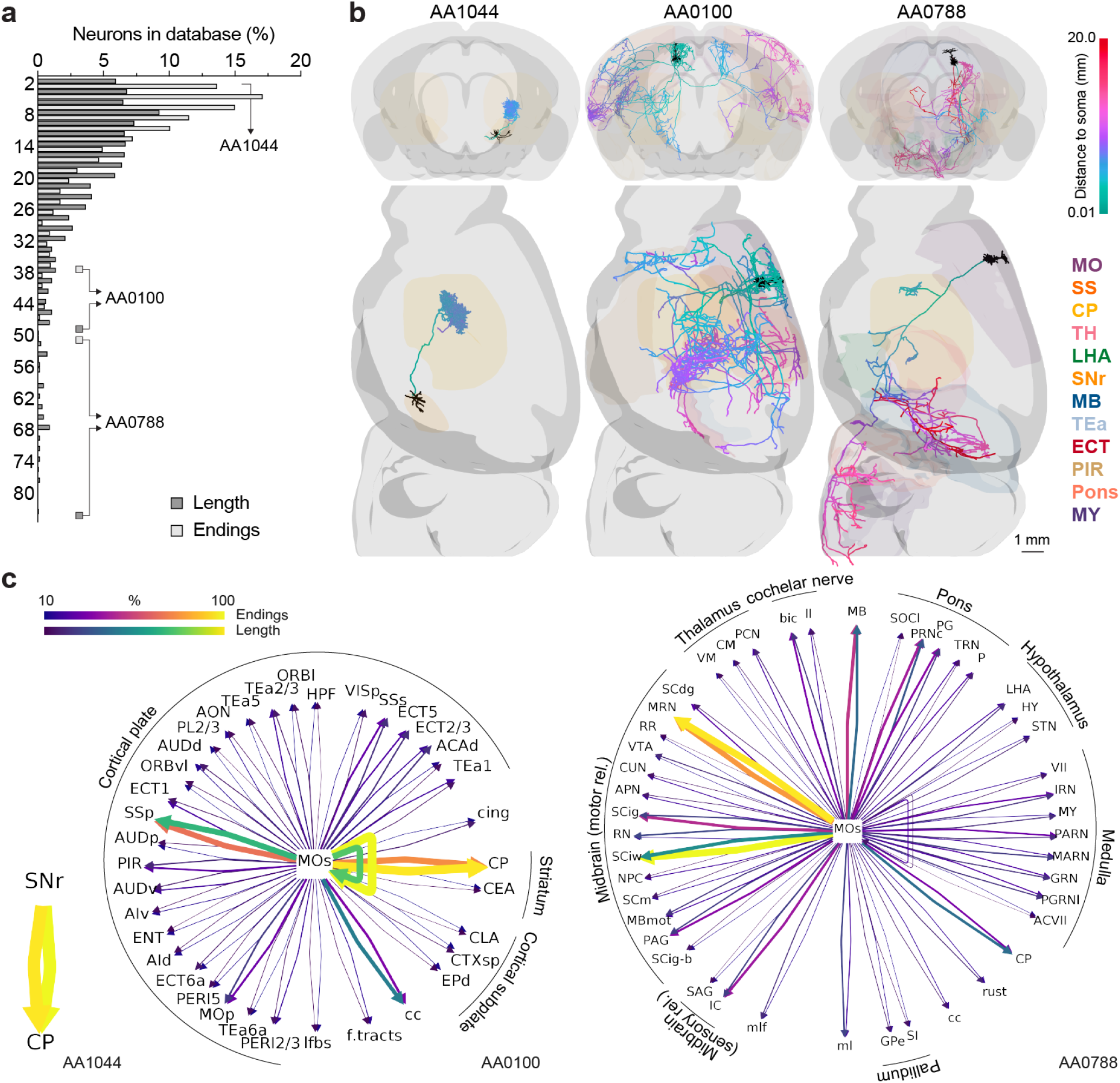
Comprehensive analytical tools enable discovery biology. *How many brain areas does a neuron connect to in the mouse brain?* The MouseLight (ML) database was programmatically parsed in SNT and the number of mid-ontology brain regions innervated by individual axons retrieved under two criteria: (normalized) cable length, and number of axonal endings at target area. (**a**) Frequency histogram of number of brain areas innervated by single cells (N=1094). (**b**) Ranked examples of axonal connectivity rendered in SNT’s *Reconstruction Viewer.* Left: SNr neuron projecting exclusively to the CP (ML id: AA1044). Its single axonal tuft can cover as much as 4% of the ipsilateral lobe of the target area. Center: SS neuron projecting to many areas in the isocortex (ML id: AA0100). Right: Pyramidal-tract neuron projecting to many areas in the isocortex, midbrain and hindbrain (ML id: AAO788). Dendrites are depicted in black and axonal arbors color-coded by “path distance to soma”, as per color ramp legend. Selected brain regions are depicted according to color-coded abbreviations (MO: Somatomotor areas; SS: Somatosensory areas; CP: Caudoputamen; TH: Thalamus; LHA: Lateral hypothalamic area; SNr: Substantia nigra, reticular part; MB: Midbrain; TEa: Temporal association areas; ECT: Ectorhinal area; PIR: Piriform area; MY: Medulla). (**c**) Connectivity diagrams for the three chosen exemplars programmatically generated in SNT’s *Graph Viewer.* In this “Ferris wheel” diagram, the neuron’s target areas are displayed around the brain area associated with the cell soma (SNr and MOs [Secondary motor area]), with connecting edges indicating projection strength, and self-connecting edges depicting local innervation. Here, edges were scaled and color-coded according to the two criteria used in a), as per color ramp legend. This representation can also be extended to cell populations (**Sup. Fig 7**). When generating such diagrams, SNT automatically sorts target areas by projection strength and groups them by parental ontology (labeled in external arcs). As a reference, the total axonal length of each cell is: 28.598cm (AA1044), 44.649cm (AA0100), 18.454cm (AA0788). Abbreviations reflect Allen Mouse Brain Common Coordinate Framework^51,52^ nomenclature.

SNT provides support for generative models of artificial neurons by utilizing the neurodevelopmental simulation framework, Cx3D^12^. This not only provides capabilities for the algorithmic generation of neuronal morphologies, but enables a new direction of image-based modeling for cellular neuroscience. On the latter, we provide a proof-of-concept example, where artificial neurons are seeded in an image derived from an *in vitro* chemotaxis assay (**Sup. Video 3**). On the former, we challenged SNT’s ability to morphometrically distinguish closely-related reconstructions. First, we generated different mathematical gene-regulatory networks (GRNs)^13^ capable of controlling neural growth by regulating extension, branching, and directionality of neurites to define *in silico* morphologies. Second, we generated thousands of artificial neurons constrained by these patterns. Third, we used built-in metrics^14,15^ to statistically differentiate between these computer-generated “neuronal types”. We found we could distinguish with high confidence all of the morphological classes, including those closely related (**Sup. Information**). Altogether, these experiments demonstrate how SNT can bridge experimental and modeled data to support model evaluation for both inference and predictive modeling.

In summary, SNT is a powerful tool for tracing, proof-editing, visualization, quantification, and modeling of neuroanatomy. It is based on recent technologies, supports modern microscopy data, integrates well with the ImageJ platform, interacts with major online repositories, and synergizes with post-reconstruction analysis software, and recent data-mining frameworks^16,17^. With a large user base and thorough community-based documentation (https://imagej.net/SNT), SNT is an accessible, scalable and standardized framework for efficient quantification of neuronal morphology.

## Methods

The figures and analyses from this manuscript can be generated programmatically. See https://github.com/morphonets/SNTManuscript for details.

### Programming

SNT was programmed with Eclipse Java IDE 4.4–4.16 (Eclipse Foundation), IntelliJ IDEA 2020 (JetBrains), and Fiji’s built-in Script Editor on an Intel i7 laptop running Ubuntu 18.10–20.04.

### Cell Image Library Imagery

Analyses were performed manually from SNT’s GUI with the following modifications to the original images: CIL810^18^ (an RGB image) was converted to a multi-channel composite; CIL701^19^ (an unannotated Z-series) was converted to a time-series stack. Please refer to the original publications for details on the datasets.

### ExFISH

*Histology:* Expansion fluorescence *in situ* hybridization (ExFISH) was performed using an optimized protocol not yet published (manuscript in preparation). In short: 150μm-thick cortical slices of an heterozygous Thy-1-GFP-M^20^ adult mouse were embedded in a hydrogel using standard embedding procedures for thick tissue^21^. mRNA was detected using HCR 3.0^22^. SST probes and fluorescent hairpins were obtained from Molecular Instruments (molecularinstruments.com). Gelled sections (~2× expanded) were imaged in PBS on a commercial Zeiss Z1 lightsheet microscope. *Spot quantification:* Signal from total RNA labeling was segmented using Labkit^23^. Individual cells were then masked using watershed filtering and labeling of connected components using MorphoLibJ^24,25^. Ill-segmented somata were manually eliminated with the aid of BAR tools^26^. Spot density (no. of spots per cell) of SST signal was determined by iteratively running *3DMaximaFinder*^27,28^ at locations of each connected component. It should be noted that this approach is rather elementary: it was designed as a proof-of-principle image processing routine that can be performed mid-way through a tracing session using accessible ImageJ tools.

### MouseLight Single-cell Connectivity

MouseLight (ML) database was programmatically parsed to obtain the number of brain areas (Allen Mouse Common Coordinate Framework (CCF) compartments) associated with individual axonal arbors using two criteria: 1) normalized cable length and 2) number of axonal endings at target area). Only CCF compartments of ontology depth 7 with public meshes available were considered. For 1) axonal length within an anatomical compartment was measured by taking all nodes within the compartment and summing the distances to their parent nodes. To exclude enpassant axons, a cell was considered to be associated with the compartment if such length would be at least 5% of the compartment’s bounding-box diagonal. For 2) only neurons with at least two end-points were considered. Selected examples in Fig. 1 were chosen by sorting cells by number of associated areas, and selecting those with the largest axonal cable length (AA1044: 28.598cm, AA0100: 44.649cm, AA0788: 18.454cm).

### Tracing and Path Fitting Benchmarks

Tracing benchmarks and fitting procedures were performed programmatically and can be reproduced using the scripts available at https://github.com/morphonets/SNTmanuscript. DIADEM scores^29^ were computed with default thresholds and retrieved in “post-DIADEM competition” mode. For degradation of traces (Fig. S2), each node in the reconstruction was displaced to a random position within a 1μm neighborhood around each axis.

### Synthetic Morphologies

*Chemoatraction assay (Video S3):* Code accessible from github.com/morphonets/ SNTManuscript. *GRNs (Fig. S6):* The code for generating GRNs is available at github.com/morphonets/cx3d/, and the five GRNs used in this study are made available at github.com/morphonets/SNTManuscript (together with remaining analysis scripts). Tools for inspecting GRNs are available at github.com/brevis-us/grneat. Morphometric analysis: Default metrics provided by SNT (41 as of this writing^1^) were retrieved for all artificial neurons. Data was normalized and analyzed using PCA *(Principal Component Analysis),* t-SNE^30^ *(t-Stochastic Neighbor Embedding),* and UMAP^31^ *(Uniform Manifold Approximation and Projection).* Group comparisons on principal components, t-SNE features, and UMAP components were performed using two-sample Kolmogorov-Smirnov tests adapted for multivariate data. p-values were combined using Fisher’s combined probability test. Density maps and examplars: soma-aligned cells were skeletonized, and their skeletons projected into the XY plane using SNT’s core functionality. Binary masks of skeletons were then summed up and resulting image normalized to the number of cells. Exemplars were chosen from a random pool of 10 cells.

## Code Availability

SNT source code is available at github.com/morphonets/SNT. The source code for the figures and analyses described in this manuscript is available at github.com/morphonets/SNTManuscript. Both are released under the GNU General Public License v3.0.

## Acknowledgments

We are extremely thankful to Jayaram Chandrashekar, Albert Cardona and Pavel Tomancak for valuable input. We thank the community of users and contributors of Simple Neurite Tracer, the developers of Scijava, pyimagej, and remaining open source libraries required by SNT, and everyone who helped test the software. W. Rasband, C. Rueden and the ImageJ community for developing and maintaining ImageJ, and Marton Rozsa and Judith Baka for critical reading of the manuscript. We thank the reviewers for constructive feedback that improved SNT. Special thanks to all the labs, teams, institutions and initiatives that facilitate public sharing of neuronal data, including 3D InsectBrain, BigNeuron (Allen Institute for Brain Science), Blue Brain, Cell Image Library, FishAtlas (MPG), FlyCircuit, FlyLight, Insect Brain Database, MouseLight, NeuroMorpho, OpenWorm, and Virtual Fly Brain.

This work was funded by the Howard Hughes Medical Institute. U.G. was funded by the Center of Advanced Systems Understanding (CASUS) which is financed by Germany’s Federal Ministry of Education and Research (BMBF) and by the Saxon Ministry for Science, Culture and Tourism (SMWK) with tax funds on the basis of the budget approved by the Saxon State Parliament.

## Author contributions

T.A.F. conceived and supervised the project. T.A.F, C.A. wrote core SNT; K.I.S.H, U.G. implemented Cx3D/sciview integration. K.I.S.H, designed and ran simulations. M.E. performed ExFISH experiment. T.A.F. analyzed the data. T.A.F., K.I.S.H., wrote the paper.

## Supplementary Information

### Overview

SNT is an open-source (GPLv3) program written in Java and distributed with the Fiji^2^ distribution of ImageJ^3^. The source code is public and managed by git (https://github.com/morphonets/SNT). SNT stems from the rewrite of Simple Neurite Tracer^4^’s basecode following Scijava^9^ design principles and with scripting abilities inspired by several powerful opensource software for neuroanatomy, namely: L-measure^14^, nat(verse)^16^, TREES^32^, btmorph^33^, and NeuroM^34^. A special effort was put into backwards compatibility, so that SNT could supersede Simple Neurite Tracer, and remain compatible with existing Fiji plugins for neuronatomy, namely *Sholl Analysis^15^.* Community-based user documentation is hosted at https://imagej.net/SNT.

SNT can run in headless environments, is fully scriptable and itself extensible. It inter-operates with other core components of the ImageJ ecosystem namely TrakEM2^35^, IJ ops^36^, and scenery^5^. SNT functionality can be extended using ImageJ2 commands^10^ or executable scripts, allowing both experienced developers and scientists with beginner-level programming experience to customize SNT. Several scripting templates are provided in Fiji’s Script Editor and a built-in discovery mechanism automatically registers user scripts in SNT’s user interface, as detailed in https://imagej.net/SNT/Scripting. Native Python is supported through pyimagej^37^. Scripting tutorials in the form of Jupyter notebooks are provided at github.com/morphonets/SNT/tree/master/notebooks.

### Requirements

An up-to-date Fiji installation running Java 8 or newer. Access to “Tubular Geodesics”^38^ segmentation requires installation of external binaries, as described in the documentation (https://imagej.net/SNT). Discrete graphics card is recommended for sciview integration. VR support in sciview requires the OpenVR/SteamVR library^5^.

### Installation

SNT is released through a dedicated “Neuroanatomy” update site, created to streamline and foster contributions from the wider scientific comunity. sciview has not been officially released and access to its functionality currently requires subscription to a second “sciview” update site. Detailed instructions are available at https://imagej.net/SNT#Installation.

### Supported File Types

**Images**: SNT accepts any non-RGB image recognized by ImageJ, SCIFIO^39^ or Bioformats^40^ with up to five axes, including multi-channel and time-lapse sequences. **Neuronal reconstructions**: SNT recognizes all known variants of SWC^41,42^, the *de facto* standard for data sharing of neuronal morphologies, MouseLight’s JSON^7^ and *Simple Neurite Tracer’s* TRACES open formats. **3D graphics**: Wavefront OBJ (reconstruction viewer) and STL, PLY, XYZ (sciview). **Analysis**: Output of SNT analyses can be saved as CSV (tabular data); SVG, PDF, PNG (plots, histograms and diagrams); XML (diagrams); MPG (sciview animations); and NeuroML (Cx3D models).

### Supported Databases

SNT can download data directly from FlyCircuit^43^ (flycircuit.tw), InsectBrainDatabase (insectbraindb.org)^6^, MouseLight^7^ (ml-neuronbrowser.janelia.org), NeuroMorpho^1^ (neuromorpho.org/), and VirtualFlyBrain^8^ (virtualflybrain.org) databases, with ongoing support for the Max Planck Zebrafish Brain Atlas^44^ (fishatlas.neuro.mpg.de/). Data can be imported from SNT’s user interface or programmatically using its API.

### Features

#### Semi-automated Tracing

The core of SNT’s semi-automatic reconstruction remains Simple Neurite Tracer’s exploratory approach in which the path between manually placed points along the centerline of neuronal processes is computed using bidirectional A* search^3^. However, several improvements were made to this procedure, namely: 1) scriptable tracing (**Fig. S1**); 2) Support for multichannel and timelapse images (**Figs. 1, S3**); 3) refinement of centerline positioning by post-hoc fitting procedures that take into account the fluorescent signal around each traced node (**Fig. S2**); 4) Detection of signal within a local 3D neighborhood around the cursor; 5) improved synchronization mechanism of Simple Neurite Tracer’s original XY/ZY/XZ tracing views that facilitate accurate node positioning; and 76) computation of curvatures on pre-processed images. The latter allows A* to be computed on mirrored data (called a “secondary image”) in which the tube-like structures of neuronal processes have been pre-enhanced using image processing routines. For added convenience, SNT offers filtering pre-sets^45,46^ through IJ-ops^36^, and allows pre-filtered data to be imported from third-party software, to e.g., allow for processing routines not yet ported into ImageJ. Adoption of other path search algorithms such as *Tubular Geodesics^38^* is also possible through the installation of external binaries. Importantly, these features can be toggled at will during a tracing session.

**Figure S1.**
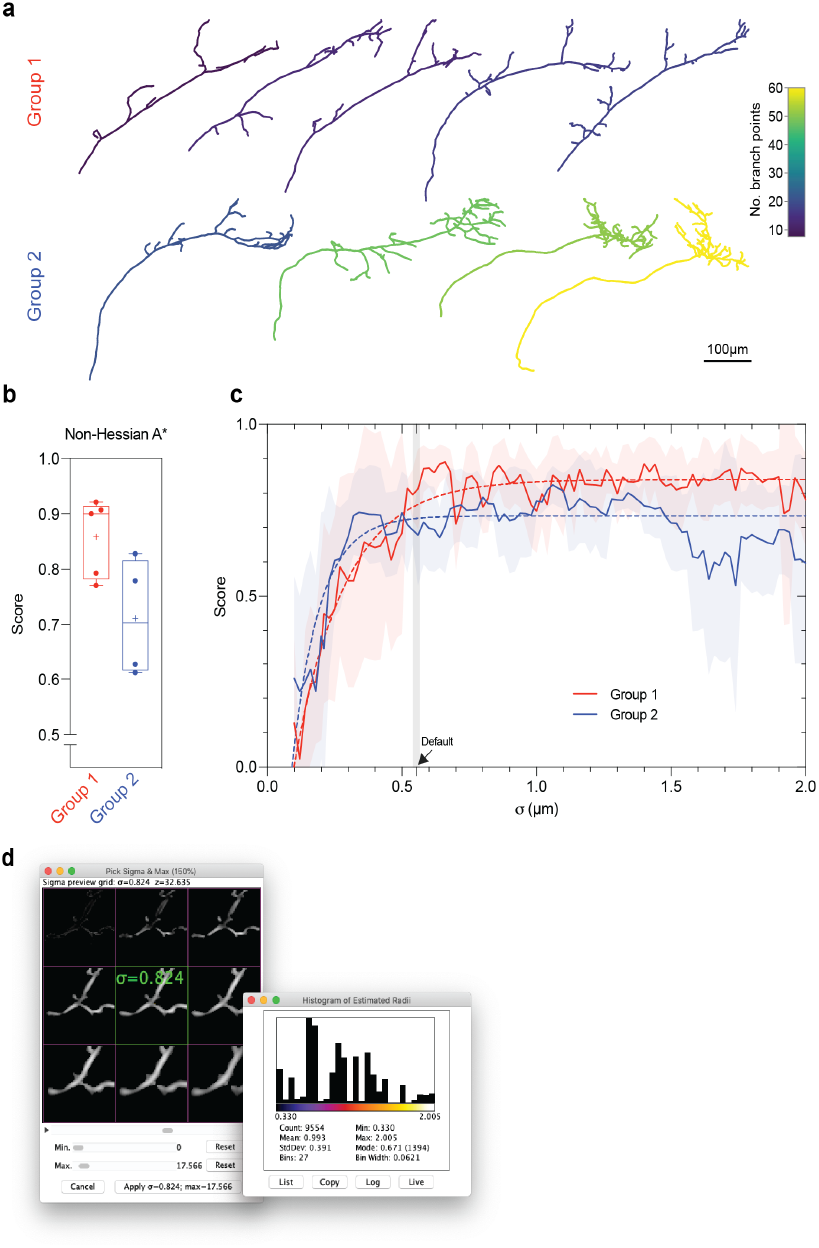
Characterization of SNT’s A* tracing in fully automated tests. **(a)** 2D renderings of axonal arbors of Drosophila olfactory projection neurons (DIADEM Challenge “OP” dataset) used in the tests, generated using SNT’s *Reconstruction Plotter* command. Cells are color-coded by number of branch points according to hue ramp. The 10 cells can be split into two complexity groups: ≤20 branches (group 1) and >20 branches (group 2). **(b)** Accuracy of automated A* without complementary Hessian analysis. Each OP cell was traced in the absence of pre-processing routines (the default in SNT) and resulting reconstruction compared to each cell’s “gold standard” using the DIADEM similarity score, in which 1 represents perfect similarity. **(c)** Effect of σ on accuracy of Hessian-based A* search. For each cell, A* search was performed on Hessian-filtered data under varying σ values. As in b), resulting reconstructions were then compared to each cell’s “gold standard”. The default value that is proposed to users in SNT’s interface is indicated (arrow). Dashed lines indicate ‘best-fit’ curves (one phase exponential). **(d)** SNT features several GUI-based tools that allow users to tune parameters during a tracing session. Here shown two controls for σ adjustment. Left: The “Hessian widget”, allowing interactive adjustments of σ at chosen image locations. Right: histogram of predicted radii, computed for the whole image volume being traced using local thickness analysis (depicted data for the OP_1 cell, the second most complex cell of group 2).

In SNT, the accuracy and performance of the automated path search can be tuned using Hessian-based analysis of curvatures, optimized to detect tubular structures of a particular size. A key parameter of this filtering operation is *σ*, the size of the Gaussian kernel used to smooth the image before detection of tube-like structures occurs^4^. In order to provide users with sensible defaults, we ran a series of simulations to assess the impact of *σ* on reconstruction accuracy. For this purpose, we chose images of Drosophila olfactory axons from the DIADEM challenge^47^, because these images share common acquisition parameters and their respective manual reconstructions (the “gold standards”) have been well characterized in benchmark studies. The topologies of these axons can be loosely divided into two complexity groups according to their number of branches (**Fig. S1a**). Since A* search can be scripted in SNT, these experiments were fully automated: For each cell, we iterated through the branches in the gold standard reconstruction and performed A* search between the voxels associated with the coordinates of the first and last node of each branch. This procedure was repeated while varying σ, with traced structures compared to the gold standard using the DIADEM metric^29^ at each run.

First, we found that in the absence of human input, the simpler cells could be traced with high accuracy in the absence of Hessian pre-processing (**Fig. S1b**, median: .90/1.00 similarity score). Second, we found that under the stringent limitations of the test, Hessian pre-processing can enhance the accuracy of the segmentation of the more complex topologies (**Fig. S1c**) but is sensitive to σ choice. It is worth noting that the default σ value —that is proposed to the user at startup upon loading of the image being traced—yielded a median DIADEM score of 0.81 for all cells combined (**Fig. S1c**). Since reconstructions associated with a score of ≥0.8 are considered acceptably similar^29,48^, SNT’s default settings —that are computed on an image-per-image basis— are reasonably determined. Given that the fine-tuning of Hessian parameters is essential for accurate results, SNT provides users with an interactive widget allowing users to adjust parameters during a tracing session, as well as commands to estimate, *a priori,* the local thickness^49,50^ of the structures to be traced (**Fig. S1d**).

SNT can also execute “path fitting” routines that automatically estimate radii and the optimal position of reconstruction nodes relative to signal (**Fig. S2a**). To expedite its usage, fitting operations are undoable, and can be perused by means of the “Fitting inspector” interface. SNT also allows users to identify and annotate portions of built-up topologies with colors and searchable tags based on custom labels, image data, or computed directly from morphometric traits (**Fig. S2b**).

#### Proof Editing and Automated Tracing

Proof-editing of automated segmentation is a major bottleneck in neuron reconstruction^*51*^. To expedite the proof-editing of traced structures, SNT features an “Edit mode” in which the topology of neuronal arbors can be edited/refined with subpixel accuracy by means of mouse clicks or keyboard shortcuts. The ability to manipulate neuron topology is essential when attempting automated reconstructions (typically involving high contrast images of simpler, unambiguous topologies). In SNT, this procedure occurs in multiple steps: the user provides a thresholded image that is skeletonized^52^. Such image is then internally converted into a graph-theoretic representation from which the reconstructed arbor is extracted. Similarly to other software packages^*53*^, this reconstruction can then be manually edited and corrected in SNT’s graphical user interface.

**Figure S2.**
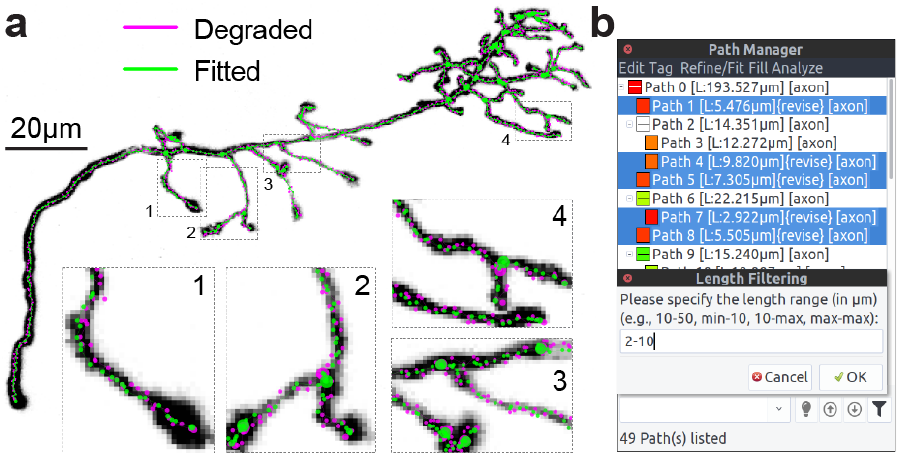
Post-hoc optimization of curvatures and proof-editing tools. **(a)** Refinement of node positioning through automated fitting. The “gold standard” reconstruction of the “OP_1” neuron (depicted as a Z-Projection) was programmatically degraded (magenta). Voxel intensities were used to correct degradation: SNT computed cross-sections along each traced path, ‘snapping’ nodes to their centroid, as depicted in insets. The same procedure can be used to estimate radii along the traced structure **(b)** The Path Manager’s search bar provides expedite commands for annotating, filtering, and selecting paths. Such operations can be based on image data, morphometric properties or user-provided tags. These are complementary to the topology-editing commands available in “Edit Mode” (not shown).

#### Multidimensional Imaging

In addition to 2D and 3D images, SNT supports multi-channel and time-lapse images, up to 5 dimensions. With time-lapse imagery, SNT allows users to associate paths associated with the same neurite across frames using a two-pronged approach: 1) tags (automated and user-based) that associate paths with neurites and frames, and 2) a matching mechanism that groups paths across frames that share a common origin. The latter can occur in a lax manner to accommodate for motion artifacts across the image (**Fig. S3a**).

**Figure S3.**
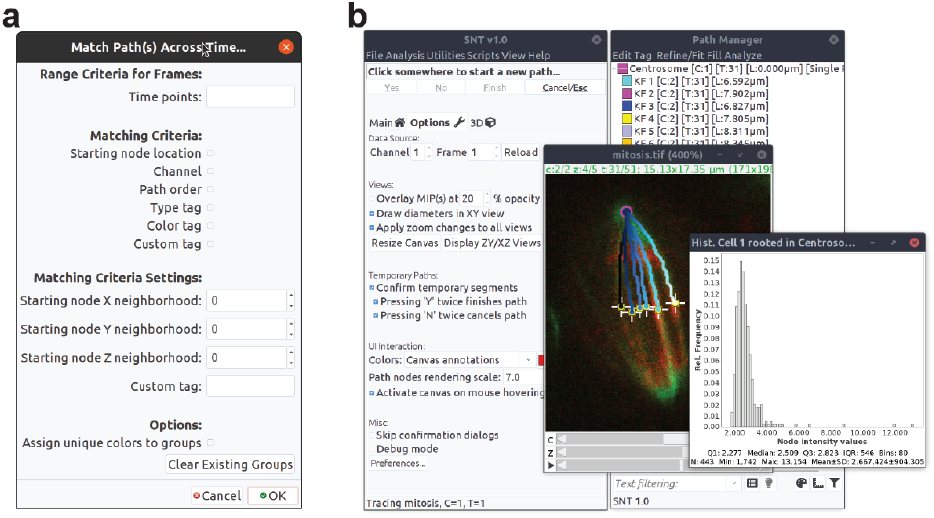
Analysis of 4D and 5D imagery. **a)** Time-lapse Analyses: Dialog of the “Match Path(s) Across Time” command showcasing the options for matching paths across time so that they remain associated with their common dendrite. The command is part of the “Time-lapse Utilities” that allow for morphometric time profiles (**Fig. 1b**). **b) Analysis of non-neuronal imagery**. K-fibers were traced during anaphase in a multi-channel 3D timelapse of dividing S2 cells^66^ (IJ1 “mitosis” 5D sample image). Traces were color-coded uniquely, according to their location along the X-axis. Histogram depicts how the voxel data underlying traced data can be easily accessed from built-in commands. Conversion of tracings into functional ImageJ ROIs is also easily accomplished, as exemplified by point ROIs (‘+’ markers) highlighting chromosome-attachment sites.

A key feature of SNT is seamless integration with ImageJ: Image processing routines can be inter-leaved with tracing tasks, and the entire suite of ImageJ plugins remains accessible during a tracing session. Special emphasis was put into allowing users to access image and reconstructed data in convenient ways. For example, reconstructed paths can be converted into functional ROIs and voxel intensities profiled along their center-lines. With multi-color fluorescence microscopy producing multi-channel images, this functionality allows users to quantify fluorescent signal along traced structures, and measure the signal from other probes in the imaged tissue (**Fig. 1c**). Although such features are of proven utility for many types of data (**Fig. S3**), we anticipate immediate utility in studies focused on neuroregeneration and neuron-glia interactions, in which the cellular environments around neurons is imaged and analyzed.

**Figure S4.**
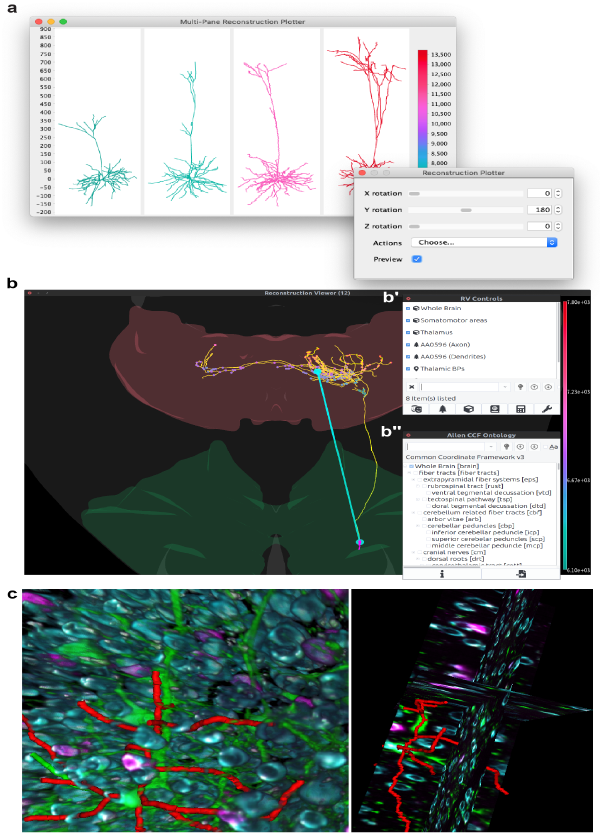
Neuroanatomy viewers. (**a**) *Reconstruction Plotter* assembles multi-panel 2D plots of reconstructed cells. Here, four dendritic arbors from the MouseLight database (IDs AA001-4), were automatically aligned, ranked and color-coded by cable length. When called from the GUI (graphical user interface), data can be transformed interactively (inset). Axis and scale bar in μm. (**b**) *Reconstruction Viewer* (RV) is a 3D visualization tool designed to handle neuroanatomy exclusively. Transverse view of the mouse brain, depicting a MouseLight pyramidal cell (ID AA0596; axon in yellow, dendrites in magenta) in the motor cortex (green). All axonal branch points within the Thalamus (red) have been identified and color-coded by depth, as per hue ramp. The vector connecting the cell soma to the centroid of thalamic branch points is highlighted. RV Controls (b’) are organized in a layout modeled after *Path Manager* (Fig. S2b) in SNT’s tracing interface. Extra functionality, such as access to Allen CCF ontologies (b”) is provided through dedicated dialogs. (**c**) Sciview integration allows SNT data to be rendered with arbitrarily large image volumes. Left: Surface visualization of the multi-channel volume described in **Fig. 1** (reconstructed dendrites in red). Right: Ortho-view of the same data (experimental feature at the time of this writing). For clarity, components of the sciview user interface (scene inspector, interactive scripting shell, Cx3D bridge, etc.) were omitted.

#### Visualization and Analysis

For simplified visualization of single-cell data SNT implements “Reconstruction Plotter”, a two-dimensional (2D) canvas for vector-based plotting of reconstructions (**Fig. S1a, S4a**). The main advantage of this type of viewer is that reconstructions can be scaled up or down to any resolution without being affected by aliasing artifacts. However, it can only render 2D data. For rendering more complex 3D data, SNT features two additional viewers: *Reconstruction Viewer* and sciview^5^.

Reconstruction Viewer (RV) is hardware accelerated, supporting both surface meshes and reconstruction files (**Fig. S4b**). The viewer can render both local and remote files on the same scene, which allows for direct loading of reconstructions from all of the supported databases and meshes for several template brains, i.e.: Drosophila (larval and adult) via Virtual Fly Brain, FlyLight, and FlyCircuit^8,43,54^; Allen Mouse Brain Common Coordinate Framework^55,56^ (CCF, adult mouse) via MouseLight database^7^; and Zebrafish via the Max Planck Zebrafish Brain Atlas^44^. In the case of the Allen Mouse Brain Atlas^56^, the full stack of anatomical ontologies is supported (**Fig. S4b”**). Reconstruction Viewer can also be used as a standalone application (i.e., in the absence of SNT’s tracing interface), allowing it to be accessed from other environments such as IPython. All Reconstruction Viewer instances can be scripted once displayed, and can be instantiated in “high-performance” mode, suitable for visualization of large amounts of data. As a proof-of-principle we used this feature to visualize the entire MouseLight database on a laptop computer without a discrete graphics card (**Sup. Video 1**). Another key feature of Reconstruction Viewer is its ability to perform geometric analyses on both meshes and reconstructions, which is key for studying the topographic organization of neurons across neuropils (**Fig. S4b**).

Sciview is a powerful SciJava-based visualization tool for volumetric and mesh-based data (**Fig. S4c**, **Sup. Video 2**). Sciview has been recently improved to support out-of-core volume rendering of images up to 9.261×10^27^ voxels^5^ (via BigDataViewer^57^), making it an appealing choice for visualization of large imagery.

SNT analytical capabilities are three-fold: 1) Analysis of imagery data already discussed; 2) Morphometric analysis of single cells in isolation and 3) Analysis of groups of cells in a common, annotated space (a reference brain/neuropil), which requires handling of neuroanatomical volumes. For morphometric analyses, SNT supports commonly used metrics^14^, graph theory based analysis, and popular quantification strategies, such as Sholl^15^ and Strahler^58–60^. In addition, SNT’s API allows for analyses based on persistent homology^61,62^ (including persistence landscapes^63^) and other ad-hoc statistical measurements complementary to those available through the user interface (**Fig. S5**).

**Figure S5.**
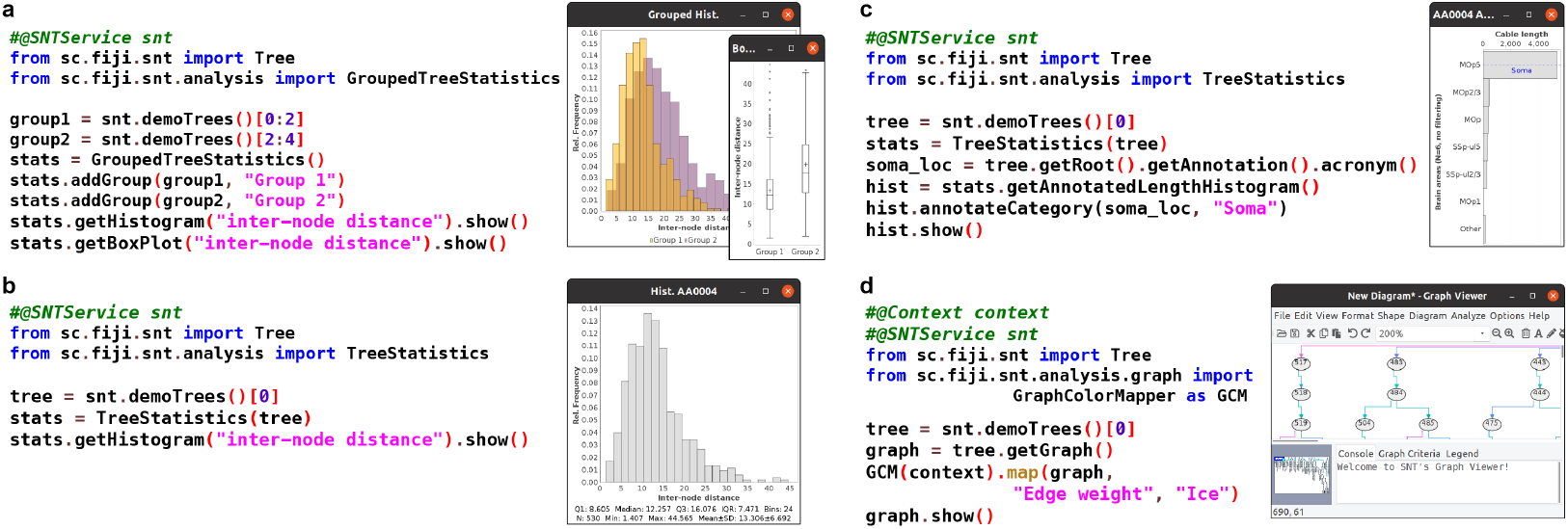
Overview of analysis API. Functional scripts (written in Python) showcasing SNT’s API (application programming interface) on dendritic arbors from the MouseLight database (cells IDs AA0001-4) that can be access offline from within SNT as “demo trees”. For each panel, the script code (as typed in Fiji’s script editor) is showed on the left. Script’s output on the right. (**a**,**b**) Statistics of morphometric traits can be extracted from groups of neurons (**a**), a single neuron (**b**), or parts thereof (not shown). Convenience methods allow data to be output in histograms, plots and tables. (**c**) Distribution of cable length across brain areas of the cell’s neuropil, in this case compartments from the Allen Mouse CCF. (**d**) Neuronal reconstructions can be converted to graphs succinctly, programmatically annotated and displayed in SNT’s interactive *Graph Viewer.* This code snippet also demonstrates how SNT interacts with the SciJava^9^ API.

#### Modeling

Cortex3D (Cx3D) is a computational modeling tool for simulating neurodevelopmental processes^12^ and has been used to define generative models of cortical circuits^64^. We integrate Cx3D with SNT through the sciview visualization package by rewriting Cx3D to grow neuronal processes with sciview’s data structures. This facilitates the use of both SNT’s and ImageJ’s functionality when designing Cx3D models. We present two important examples that benefit from the unification of quantitative neuroanatomy and morphological modeling: statistical discrimination between morphological cell types, and image-based modeling of neural morphologies. However, there are many additional possibilities enabled by this coupling, from generative testing of new quantitative measurements to data-driven model fitting.

#### Morphological Discrimination Between Cell Types

Morphological discrimination between neuronal cell types is an important aspect of cell classification. The quantitative measurements made by SNT can be used to statistically test the difference between morphological populations of cells. To test this, we generated thousands of unique artificial neurons that ‘developed’ *in silico* under similar conditions and asked if SNT metrics could resolve such morphologically-related topologies.

First we integrated a computational model of artificial gene regulatory networks (GRNs)^13^ into Cx3D for the control of cellular processes. An overview of the GRN architecture and mechanism of simulating GRN dynamics is described in detail in the previous publication^13^. Second, we generated five unique GRNs. While each of the five unique GRNs are randomly generated, all GRNs are provided with common inputs: directional cues for each 3-dimensional axis, length of the neurite segment, volume of the neurite segment, and local branch order. Additionally, all GRNs have common outputs: directional bias for each 3-dimensional axis, directional bias for previous direction of growth, directional bias for taking a random direction, a factor for increasing the segmentation of a neurite, and a factor for increasing the probability of branching. This type of input/output behavior is described extensively in previous publications of the Cx3D simulator^12,64^. The specific connectivity and number of regulatory components of the GRN were randomized. However, completely randomized GRNs have a low probability of being capable of generating viable morphologies. To address this, rejection sampling was used when generating the random GRNs by testing that each network satisfied a minimal viability criteria. The viability test was performed as follows: the GRN is simulated for 100 time-steps with constant input values, with the following criteria assessed at the first and final time-step of the viability test: (1) all behavior regulating protein concentrations should change, (2) the GRN must bifurcate enough to have reasonable growth, (3) the GRN must not over-bifurcate, (4) the GRN must trigger branching, and (5) the GRN must not overbranch. While these criteria will have an impact on the distribution of possible GRNs that are selected, the rationale for this approach is to compensate for the fact that most randomly generated GRNs are not capable of generating physically plausible neurodevelopment.

**Figure S6.**
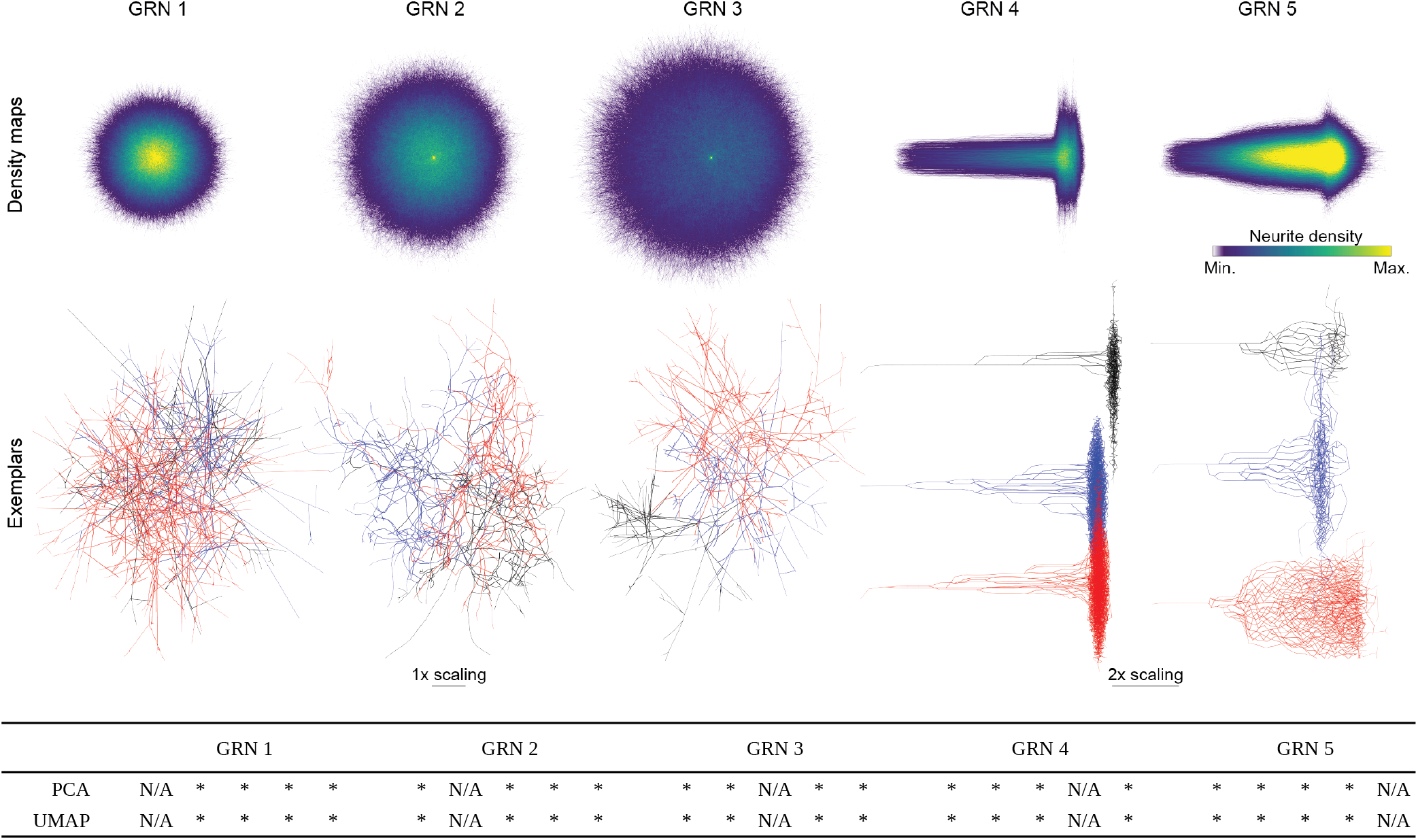
Statistical discrimination between between morphological populations of cells generated *in silico.* Top: Density maps for the five neurodevelopmental cell types artificially generated using unique GRNs, obtained by overlaying all soma-aligned cells in the group. Color coding reflects the density of neurites at a given location for the 5 groups. Middle: Representative exemplars of each group. For clarity, cells from GRNs 4 and 5 were separated by a vertical offset. Bottom: Summary of pairwise statistical measures of morphological differences between the five groups analyzed under different dimensionality reduction techniques. Principal components, t-SNE features (not shown) and UMAP components were computed from default SNT metrics and compared using a modified two-sample Kolmogorov Smirnov test. *: p<0.001; N/A: no test performed between same sample. Computed results available at https://github.com/morphonets/SNTManuscript. Cells from GRNs 4 and 5 were re-scaled in top and middle panel so that all cells could be rendered under equivalent dimensions. N=1000 cells per group.

For each GRN, we stochastically generated 1,000 artificial neurons—under the same environmental conditions— and analyzed their skeletons with SNT core metrics (**Fig. S6**). We then performed Principal Component Analysis (PCA) and statistically tested the difference between principal components with two-sample Kolmogorov-Smirnov tests combined with the Fisher combined probability test (**Methods**). We obtained statistical significance on all comparisons. All groups were similarly resolved when extending this approach to other dimensionality reduction techniques (t-SNE^30^ and UMAP^31^).

#### Image-based Modeling

By unifying SNT and Cx3D, we can leverage ImageJ within Cx3D to enable support for image-based modeling. We demonstrate a proof-of-concept simulation based upon an *in vitro* microfluidic assay designed for screening the effect of chemoattractants on neuronal growth^65^. We recreated their microfluidic circuit topology as a 3-dimensional image, and define gradients of chemoattractant using intensity values of the image_voxels, analogous to the original experiment’s Netrin assay. Neurite outgrowths follow chemical gradients of increasing concentration defined within the microfluidic circuit. We show a simulated neuron and image-based environment in **Sup. Video 3**.

The neurodevelopment model encodes a minimal artificial environment reminiscent of an *in vitro* assay. A neuron is initialized as a single soma at a randomly selected position within +/− 40 spatial units of the origin (0, 0, 0) along the X-, Y-, and Z-axes. Three extracellular morphogen gradients are established and extend for 300 spatial units along the three axes in a Gaussian distribution concentration. The simulation begins by extending an initial neurite segment from the soma. The GRNs then regulate the growth of neurites with the previously described inputs and outputs. The model proceeds by iteratively simulating physical constraints encoded in the Cx3D simulation engine and the dynamics of the GRN to grow the artificial neuron.

## Glosssary

GRNs: Artificial gene regulatory networks (GRNs) are mathematical algorithms inspired by mechanisms of biological gene regulation. GRNs can be used to model or solve problems with a strong dynamic or stochastic component
Mesh: A polygon mesh defines the shape of a three-dimensional polyhedral object. In neuronal anatomy, meshes define *neuropil* annotations, typically compartments of a reference brain atlas (e.g., the hippocampal formation in mammals, or mushroom bodies in insects)
Multi-dimensional image: An image with more than 3 dimensions (3D). Examples include fluorescent images associated with multiple fluorophores (multi-channel) and images with a time-dimension (time-lapse videos). A 3D multi-channel timelapse has 5 dimensions
Neurite: Same as neuronal process. Either an axon or a dendrite
Path: Can be defined as a sequence of branches, starting from soma or a branch point until a termination. In manual and assisted (semi-automated) tracing, neuronal arbors are traced using paths, not branches. Fitting algorithms that take into account voxel intensities can be used to refine the center-line coordinates of a path, typically to obtain more accurate curvatures. Fitting procedures can also be used to estimate the volume of the neurite(s) associated with a path
(Neuronal) morphometry: Quantification of neuronal morphology
Neuropil: Any area in the nervous system. The cellular tissue around neuronal processes
Out-of-core: Software with *out-of-core* capabilities is able to process data that is too large to fit into a computer’s main memory
Reconstruction: See *Tracing*
ROI: Region of Interest. Define specific parts of an image to be processed in image processing routines
Skeleton: A thinned version of a digitize shape (such as a neuronal reconstruction) or of a binary image
Tracing: A digital reconstruction of a neuron or neurite. The term predates computational neuroscience and reflects the manual ‘tracing’ on paper performed with camera lucida devices by early neuroanatomists
Volume rendering: A visualization technique for displaying image volumes (3D images) directly as 3D objects

## Supplementary Videos

**Video S1 |** Example of a programmatic animation: “Cumulative” rendering of the complete MouseLight database in *Reconstruction Viewer.* Animation, was generated on a laptop computer lacking a dedicated GPU, using a single script that downloaded, measured, and rendered each cell. See https://github.com/morphonets/SNTManuscript for details. Note that the number of cells in the database has meanwhile surpassed those rendered.

**Video S2 |** Showcase sexample of scivew capabilities, in which segmentation and volumetric data are rendered in the same scene. All data was loaded from the *Cremi* challenge (https://www.cremi.org) sample dataset “A”, with the ten largest volumes (by voxel count) shown in random colors. In addition, half the volume of the EM raw data is shown as a semi-transparent direct volume rendering.

**Video S3 |** Example of image-based modeling using Cx3D and a 3D volumetric image defining a microfluidic circuit designed to assess neurite outgrowth in response to Netrin-1 and and Slit2 ^65^. The simulation shows a single cell (magenta) with chemotaxis and branching preference for a steady-state chemical gradient (low concentration: purple; high concentration: yellow).

**Figure S7.**
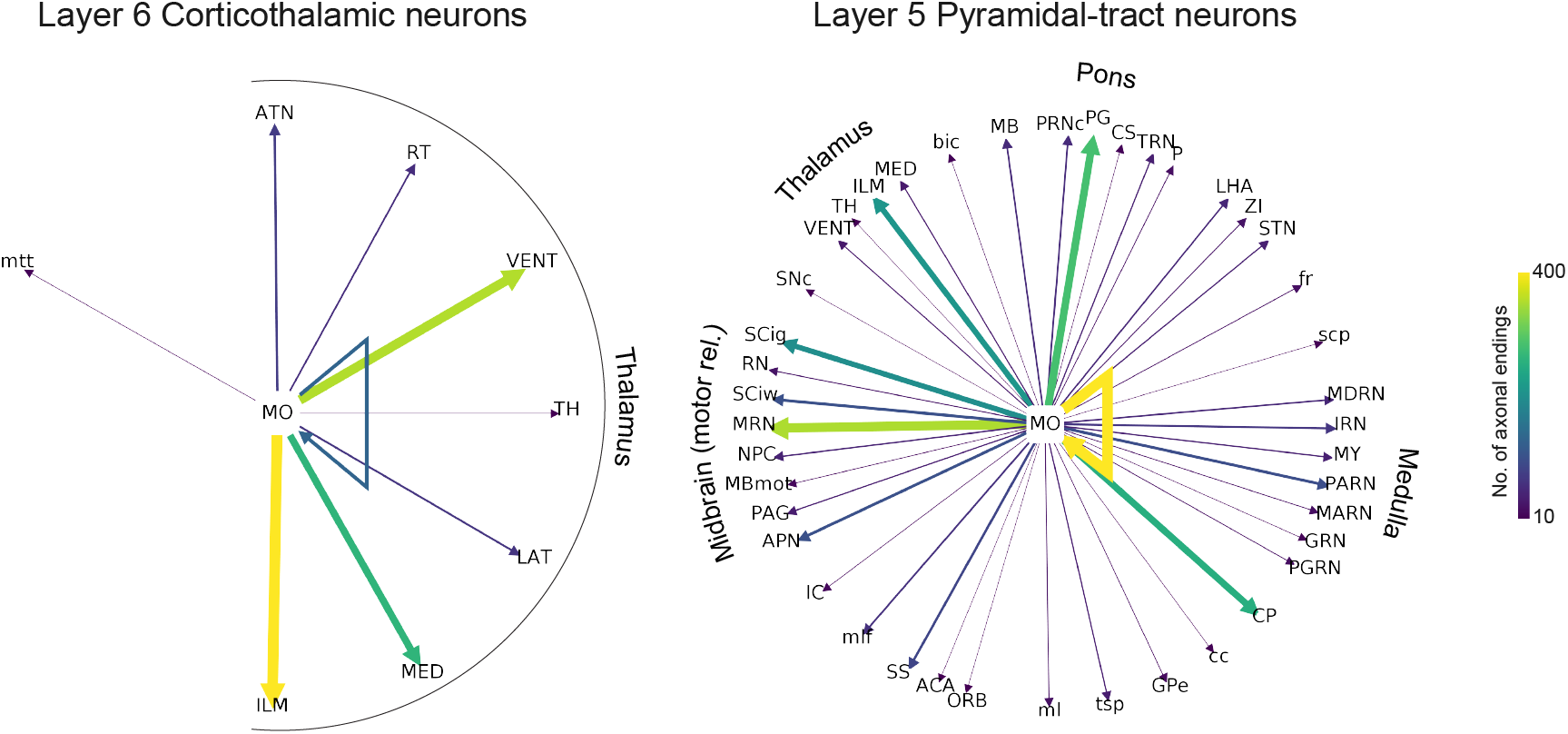
Connectivity “Ferris Wheel” diagrams for two cell populations in Layer 5/6 of the secondary motor area in the mouse brain. The two populations are described in detail in Fig. 6 of Winnubst et al^7^. For simplicity, only brain areas at midontology level (depth=6) and one morphometric criteria (no. of axonal endings at target location) were considered. The cells were retrieved from the MouseLight database and, as in **Fig. 2**, diagrams were programmatically generated (see https://github.com/morphonets/SNTManuscript for details). No of cells per group: 20 (corticothalamic); 14 (pyramidal-tract).

## Footnotes

1 A subset of all available metrics: *Average branch length, Average contraction, Average fractal dimension, Average fragmentation, Average partition asymmetry, Average remote bif. angle, Cable length, Depth, Height, Highest path order, Horton-Strahler bifurcation ratio, Horton-Strahler number, Length of inner branches (sum), Length of primary branches (sum), Length of terminal branches (sum), Mean radius, No. of branch points, No. of branches, No. of inner branches, No. of nodes, No. of primary branches, No. of terminal branches, No. of tips, Width;* [Sholl-based metrics]: *Centroid, Centroid radius, Decay, Degree of polynomial fit, Enclosing radius, Intercept, Kurtosis, Max, Max (fitted), Max (fitted) radius, Mean, Median, No. maxima, No. secondary maxima, Skeweness, Sum, Variance.* Refer to user documentation for details.

